# Stimulation related artifacts and a multipurpose template-based offline removal solution for a novel sensing-enabled deep brain stimulation device

**DOI:** 10.1101/2021.09.10.457001

**Authors:** Lauren H Hammer, Ryan B Kochanski, Philip A Starr, Simon Little

## Abstract

**Background:** The Medtronic “Percept” is the first FDA approved deep brain stimulation (DBS) device with sensing capabilities during active stimulation. Its real-world signal recording properties have yet to be fully described.

**Objective:** This study details sources of artifact (and potential mitigations) in local field potential (LFP) signals collected by the Percept, and assesses the potential impact of artifact on the future development of adaptive DBS (aDBS) using this device.

**Methods:** LFP signals were collected from seven subjects in both experimental and clinical settings. The presence of artifacts and their effect on the spectral content of neural signals were evaluated in both the stimulation ON and OFF states using three distinct offline artifact removal techniques.

**Results:** Template subtraction successfully removed multiple sources of artifact, including 1) electrocardiogram (ECG), 2) non-physiologic polyphasic artifacts, and 3) ramping related artifacts seen when changing stimulation amplitudes. ECG removal from stimulation ON (at 0 mA) signals recovered the spectral shape seen when OFF stimulation (averaged difference in normalized power in theta, alpha, and beta bands ≤ 3.5%). ECG removal using singular value decomposition was similarly successful, though required subjective researcher input. QRS interpolation produced similar recovery of beta-band signal, but resulted in residual low-frequency artifact.

**Conclusions:** Artifacts present when stimulation is enabled notably affected the spectral properties of sensed signals using the Percept. Multiple discrete artifacts could be successfully removed offline using an automated template subtraction method. The presence of unrejected artifact likely influences online power estimates, with the potential to affect aDBS algorithm performance.

## Introduction

Deep brain stimulation (DBS) is a treatment for patients with numerous neurologic conditions, including Parkinson’s disease [1–4], essential tremor [5–7], and dystonia [8–11], for whom pharmacologic therapy is inadequate. Standard-of-care currently delivers DBS in a continuous manner, without automated feedback to adjust therapy according to changing motor signs. Recent work has focused on the development of adaptive DBS (aDBS), where stimulation is modulated in response to a biomarker of the patient’s clinical state [12]. Neurophysiologic biomarkers, such as signal properties of subcortical local field potentials (LFPs) recorded from the DBS lead itself, are frequently proposed as feedback signals for aDBS systems [13,14]. For example, beta range (13-30 Hz) oscillations recorded from the subthalamic nucleus (STN) have correlated with symptoms of Parkinson’s disease [13], and beta-band power was successfully implemented as a control signal for laboratory-based implementations of aDBS [15–17]. Similar paradigms were trialed in cervical dystonia using lower frequency bands (4-12 Hz) recorded from the globus pallidus (GP) [18]. Successful application of aDBS using subcortical LFP biomarkers therefore relies on accurate sensing of neural signals, particularly within frequency bands of interest.

Initial studies investigating aDBS utilized externalized leads connected to benchtop amplifiers, which allowed LFPs to be recorded during active stimulation with a high signal-to-noise ratio [15,19]. Subsequent work has focused on implementing such algorithms within fully implanted devices. Medtronic’s first-generation investigational DBS internal pulse generator (IPG) with sensing capabilities, the Activa PC+S, was successfully used in investigational aDBS systems for Parkinson’s disease [17] and essential tremor [20], though other studies described difficulty sensing subcortical LFPs during active stimulation because of artifact [21–23]. Sources of artifact included stimulation artifact [21,23], electrocardiogram (ECG) [21,24,25], and clock noise [21]. Offline post-processing attempts to remove stimulation artifact were proposed, but were generally unable to fully remove it [26]. Medtronic’s next generation investigational device, the Summit RC+S, included design specification changes to address issues with artifact, including sense blanking, implementation of a fully differential amplifier, active recharge, and improved connector seals [27]. However, the RC+S remains limited to a small number of centers and only in the research setting [28].

Recently, the Medtronic Percept PC became the first FDA approved device with the capability to sense LFPs during stimulation [29]. Preliminary reports have highlighted the presence of ECG artifacts, particularly when stimulation is turned ON [31,32] (ON is a device status setting, and can be configured to deliver no current if amplitude is set to 0 mA). Other sources of artifact included movement artifact and those caused by turning stimulation OFF or ON. Artifacts caused by ramping (changing stimulation amplitude), as has been seen with the RC+S [27], have not been previously characterized. Two previous studies (pre-print) have noted the presence of ON-stimulation artifact [31,32] and proposed methods for artifact removal (focusing on ECG), though there has not been a direct comparison of multiple methods. Nor has there been a comprehensive comparison of the LFP signal properties recorded when stimulation is ON versus OFF after artifact removal. Given the Percept offers multiple LFP sensing functionalities (Table 1), future experimental designs using the Percept will rely on whether direct comparisons can be made between signals recorded by different methods. The objective of this work was to therefore assess the differences in signal properties recorded when stimulation was ON versus OFF, including characterizing the presence of artifact (such as ECG and those provoked by changes in stimulation amplitude), assessing the effects of artifact on spectral content, and comparing multiple techniques for artifact removal.

**Table 1:** Percept LFP Recording Functionalities.

| **Sensing Method** | **Recording Duration** | **Environment** | **Bipolar Channels Used** | **Sensing During Stimulation?** | **Time-Domain Data?** | **On-Board Calculated Spectral Data?** |
| --- | --- | --- | --- | --- | --- | --- |
| Signal Quality Check - Sense Channel Test | ~20 s | In clinic / lab | Stimulation-compatible* | No | Yes | Yes, full spectrum of the 20 s data set |
| Signal Quality Check - Calibration Test | ~20 s | In clinic / lab | Single (sandwiching stimulation cathode)** | Yes (0 mA stimulation only) | Yes | No |
| LFP Montage | ~20 s | In clinic / lab | All*** | No | Yes | Yes, full spectrum of the 20 s data set |
| Indefinite Streaming | Continuous streaming | In clinic / lab | Stimulation-compatible* | No | Yes | No |
| Brain Sense Streaming | Continuous streaming | In clinic / lab | Single (sandwiching stimulation cathode) | Yes | Yes | Yes, pre-defined 5-Hz narrow band power, updated every 10 min |
| Brain Sense Events | ~30 s | At home | Single (sandwiching stimulation cathode) | Yes | No | Yes, full spectrum of the 30 s data set |
| Brain Sense Timeline | Continuous streaming | At home | Single (sandwiching stimulation cathode) | Yes | No | Yes, pre-defined 5-Hz narrow band power, updated every 10 min |

\* Stimulation-compatible bipolar channels include only bipolar configurations that “sandwich” the cathode in monopolar or double monopolar stimulation settings using the middle contacts (i.e. 0-2, 0-3, 1-3, 8-10, 8-11, and 9-11).
\*\* Consecutively assesses configurations compatible with sensing during single monopolar stimulation (i.e. 0-2, 1-3, 8-10, and 9-11).
\*\*\* Consecutively assesses all configurations compatible with stimulation\*, and then all configurations not compatible with stimulation (i.e. bipolar recording pairs are adjacent contacts)

## Methods

### Subjects

Seven subjects participated in this study (Table 2) after informed consent to protocols approved by the local institutional review board. Each had a Percept IPG connected to Medtronic 3389 leads, implanted in either the STN or GP. Subject Sub05 also had a subdural cortical paddle (Medtronic Resume II 3587A25) implanted over the left primary motor cortex (M1) from a previous investigational protocol [21].

**Table 2:** Subject Demographics.

| Subject | Age / Sex | Diagnosis | Recording Setting | IPG Site | Lead Target | Stimulation Contacts (Sensing Contacts) | Type of Artifact Identified by Percept |
| --- | --- | --- | --- | --- | --- | --- | --- |
| Sub01 | 46 / Male | Holmes tremor | Experimental | Left chest | Left GP | C+2- (1 and 3) | None |
| Sub02 | 73 / Male | Parkinson's disease | Experimental | Right chest | Left STN | C+1- (0 and 2) | None |
|  |  |  |  |  | Right STN | C+9- (8 and 10) | None |
| Sub03 | 80 / Female | Parkinson's disease | Experimental + Clinical | Left chest | Right GP | C+9- (8 and 10) | SQC |
| Sub04 | 66 / Female | Parkinson's disease | Experimental | Left chest | Left GP | C+1- (0 and 2) | CT |
|  |  |  |  |  | Right GP | C+9- (8 and 10) | SQC |
| Sub05 | 46 / Male | Parkinson's disease | Clinical | Left chest | Left STN | C+1- (0 and 2) | CT |
|  |  |  |  |  | Left M1 | n/a (1 and 3) | SQC |
| Sub06 | 70 / Male | Parkinson's disease | Clinical | Right chest | Left GP | C+1- (0 and 2) | None |
|  |  |  |  |  | Right GP | C+9- (8 and 10) | None |
| Sub07 | 72 / Male | Parkinson's disease | Clinical | Left chest | Left GP | C+1- (0 and 2) | None |

SQC, signal quality check (assessment when stimulation is off); CT, calibration test (assessment when stimulation is on with amplitude at 0 mA, (ON-0mA))

### Data Acquisition

LFP signals were collected at 250 Hz from all subjects using the “Brain Sense Streaming” mode (Table 1). DBS programming for each subject was configured to monopolar stimulation at one of the two inner contacts (contacts 1/2 or 9/10), and LFPs were sensed using the two contacts adjacent to the cathode in a bipolar configuration (Table 2).

Four subjects (Sub01-Sub04) participated in a defined experimental paradigm. Time series data were collected from subjects while at rest, with stimulation either (1) OFF or (2) ON with amplitude equal to 0 mA (ON-0mA state). Two subjects had unilateral implants and two subjects had bilateral implants, resulting in LFP recordings from six individual hemispheres. Five repetitions were collected for each subject in both stimulation states (only four repetitions of ON-0mA for Sub02). The order of stimulation conditions was randomized.

LFPs were also collected from four subjects (Sub03, and Sub05-Sub07) during clinical care and analyzed offline. Time series data were collected simultaneously with the stimulation amplitude (which was updated every 0.5 s). For Sub07, 20 s of OFF-stimulation time-series data were also collected using the “LFP Montage” (Table 1).

### Offline Artifact Removal

Time series data were first low-pass filtered using a seventh-order Butterworth filter with a cutoff frequency of 100 Hz. Three methods of ECG removal were assessed: a new template subtraction pipeline, singular value decomposition (SVD), and QRS interpolation. The template subtraction technique was also implemented to remove non-ECG repetitive artifacts.

### Template Subtraction Pipeline

Template subtraction [33] consisted of a four-step process (Fig. 1). The process is described below in detail for ECG, along with modifications for use with other repetitive non-ECG artifacts.

**Fig. 1:**
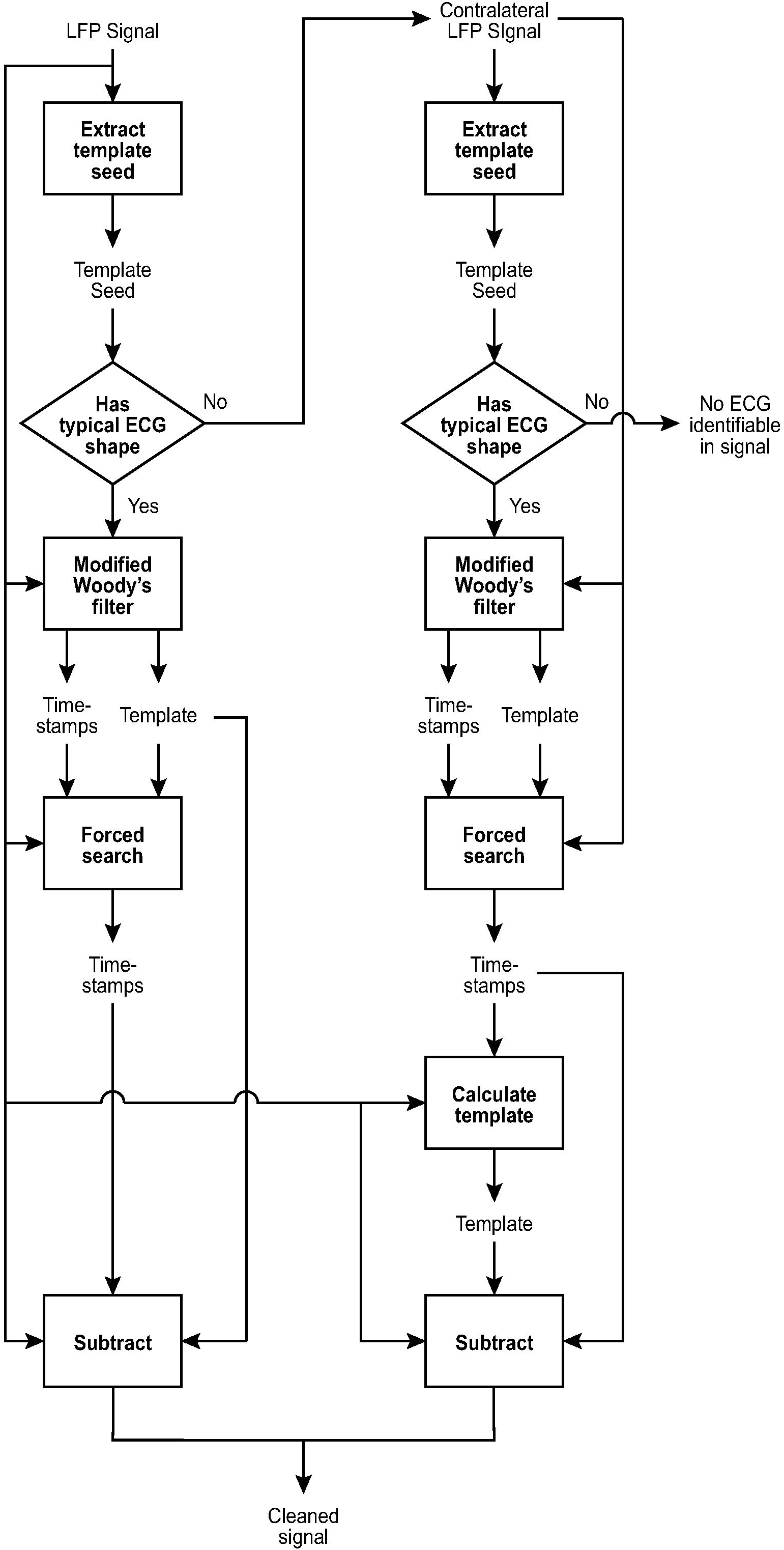
Summary of template subtraction pipeline for ECG removal. Template subtraction consisted of a four-step process, including (1) identification of a template seed (estimate of the artifact waveform), (2) use of a modified Woody’s adaptive filter to identify locations of artifact and recursively update the template shape, (3) a forced search for artifact locations potentially missed by the Woody’s filter, and (4) subtraction of the finalized template shape at identified timestamps.

1. Template Seed Identification: For ECG removal, template seed identification was automated. Each possible 760 ms epoch within the first 5 s of the signal was evaluated for possible ECG waveforms (760 ms epoch was based on the upper limits of normal for PR and QT intervals of 200 ms and 460 ms respectively, along with a 100 ms buffer). Each epoch was used as the “known waveform” in a matched filter on the remaining signal, where peaks of the matched filter output represented timestamps of greatest correlation with the epoch’s waveform. The epoch with the highest average peak filter output was identified (with a constraint that the peaks be at least 0.5 s apart, equivalent to a heart rate of 120 bpm). The template seed was set as the average of these waveform matches.
2. Modified Woody’s Adaptive Filter: Once a template seed was identified, a Woody’s adaptive filter [34] was used to identify timestamps of maximum correlation with the template and iteratively update the template shape. The Woody’s filter was implemented using a matched filter to identify the timestamps of maximum correlation, with the constraint that the timestamps be at least 0.5 s apart (equivalent to a heart rate of 120 bpm). To minimize the false positive rate, the matched filter output was also required to exceed a threshold (97.5th percentile of filter output for the entire signal) to be included.
3. Forced Search for Missed Artifact Identification: Threshold use with the Woody’s filter resulted in decreased sensitivity (but higher specificity) of artifact detection. Potential timestamps of missed artifact detection were identified by finding epochs where the inter-timestamp duration exceeded 1.5 times the mode. Within each identified epoch, the number of missed artifacts was estimated (N). A matched filter was used to find N timestamps with the greatest correlation to the Woody’s filter’s output template.
4. Subtraction of Artifact Template: Once all artifact timestamps were identified, a cleaned signal was produced by subtracting the Woody’s filter output template from the signal at the determined timestamps. If the duration between timestamps was smaller than the length of the template, the template was trimmed to prevent overlap and redundancy of template subtraction.

For signals where the automated template seed identification did not produce ECG artifact, the subtracted ECG template was produced by averaging epochs at the artifact timestamps identified in the contralateral hemisphere. If neither hemisphere produced an automated ECG template seed, and no ECG could be observed by visually inspecting the signals, the signals were considered to be absent of ECG. For experimental conditions with multiple trials (e.g. experiments comparing the signal properties when stimulation is OFF versus ON-0mA), the template seeds and the final templates subtracted from the signals were averaged across all trials within the same subject and experimental condition.

For implementation of template subtraction with non-ECG artifact, the template seed was manually selected as an epoch of signal including artifact. The required minimum time between peaks of matched filter output was ninety percent of a manual estimate of the typical duration between artifact incidences.

### Singular Value Decomposition (SVD)

SVD was implemented as previously described for ECG removal [25,32]. In brief -- epochs of M samples (corresponding to 760 ms, or trimmed as above if epochs overlapped) were extracted at timestamps of ECG incidence (identified by the same technique used with template subtraction). For experimental conditions with multiple trials, ECG epochs were grouped within each combination of subject and experimental condition, producing a MxP matrix (where P is the number of ECG artifacts). SVD of this matrix produced a set of projections equivalent to projections onto principal component eigenvectors. Eigenvector projections that extracted signals consistent with ECG morphology were identified by visual inspection and one or more projections were selected as artifact-related. Artifact-related projections were then used to reconstruct ECG estimates at each timestamp, which were subsequently subtracted.

### QRS Interpolation

QRS interpolation was a modified version of an online repository script for processing signals collected from the Percept [31]. This method identifies the location of QRS complexes, and the artifact is removed by interpolation, where the QRS complex is replaced by the signal immediately preceding and immediately following the identified QRS. This method was implemented as published in the repository, except QRS complexes were instead identified using 100 ms epochs within ECG artifacts identified by the template subtraction method for consistency.

### Offline Spectral Analysis and Power Normalization

For subjects Sub01-Sub05, the power spectra of signals collected in the OFF and ON-0mA state were calculated, both before and after artifact removal. For subjects with bilateral DBS implants, each hemisphere was assessed. The power spectra of LFP data were calculated using the Welch method, with a 1 s window and 50% overlap. For Sub01-Sub04, the integrated power within three spectral bands (theta: 4-7.5 Hz, alpha 8-13 Hz, and beta 13.5-30 Hz) was calculated for each trial. The average percent difference of power between cleaned ON-0mA and OFF-stimulation signals were calculated for each power band and ECG removal technique, and then averaged across subjects. To account for possible broad spectral scaling differences between OFF and ON-0mA sensing conditions (which affect absolute power estimates but not shape of the power spectrum), spectral power was normalized by integrated gamma power (30-125 Hz), given the minimal gamma content of ECG artifact.

To assess how artifacts might impact embedded estimates of spectral power (“Brain Sense” mode), we calculated estimates of narrow band (5 Hz-wide) power centered on two frequencies in the alpha and beta ranges. Power estimates were updated every 3 s, where the Welch method was implemented on the preceding 3 s epoch, using a 1 s window and 50% overlap.

## Results

### ECG Artifact

LFP signals recorded by the Percept in Sub01-Sub04 demonstrated notably different power spectra when collected with stimulation OFF versus ON-0mA (Fig. 2), and were primarily caused by the presence of ECG artifact (Fig. 3A). For two hemispheres (Sub02-R and Sub 04-R), ECG artifact was not visually apparent in ON-0mA signals, and was only extracted by averaging epochs at timestamps identified from the contralateral hemisphere. No ECG artifact could be identified by visual inspection or by automated template seed generation for any subject when recorded in the OFF-stimulation state.

**Fig. 2:**
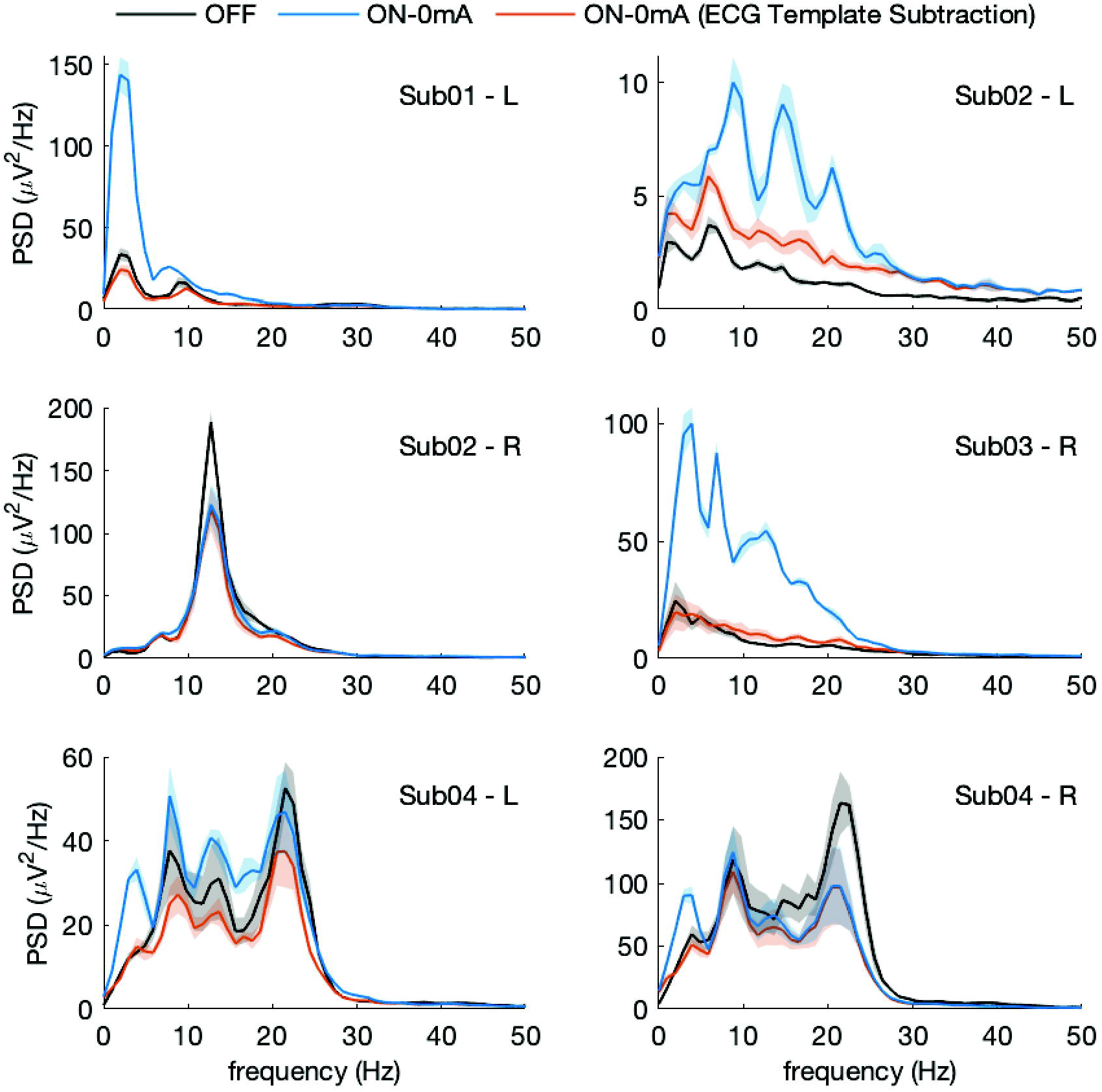
LFP spectral content when stimulation is OFF versus ON-0mA. Turning on stimulation (though not delivering any current) resulted in significant change in the LFP power spectrum. Removal of ECG artifact from ON-0mA signals using template subtraction resulted in the recovery of the scaled spectral shape seen with signals collected when stimulation was OFF. However, there was not a consistent scaling factor between subjects/hemispheres. Shaded region represents standard error. L, left hemisphere. R, right hemisphere. PSD, power spectral density.

**Fig. 3:**
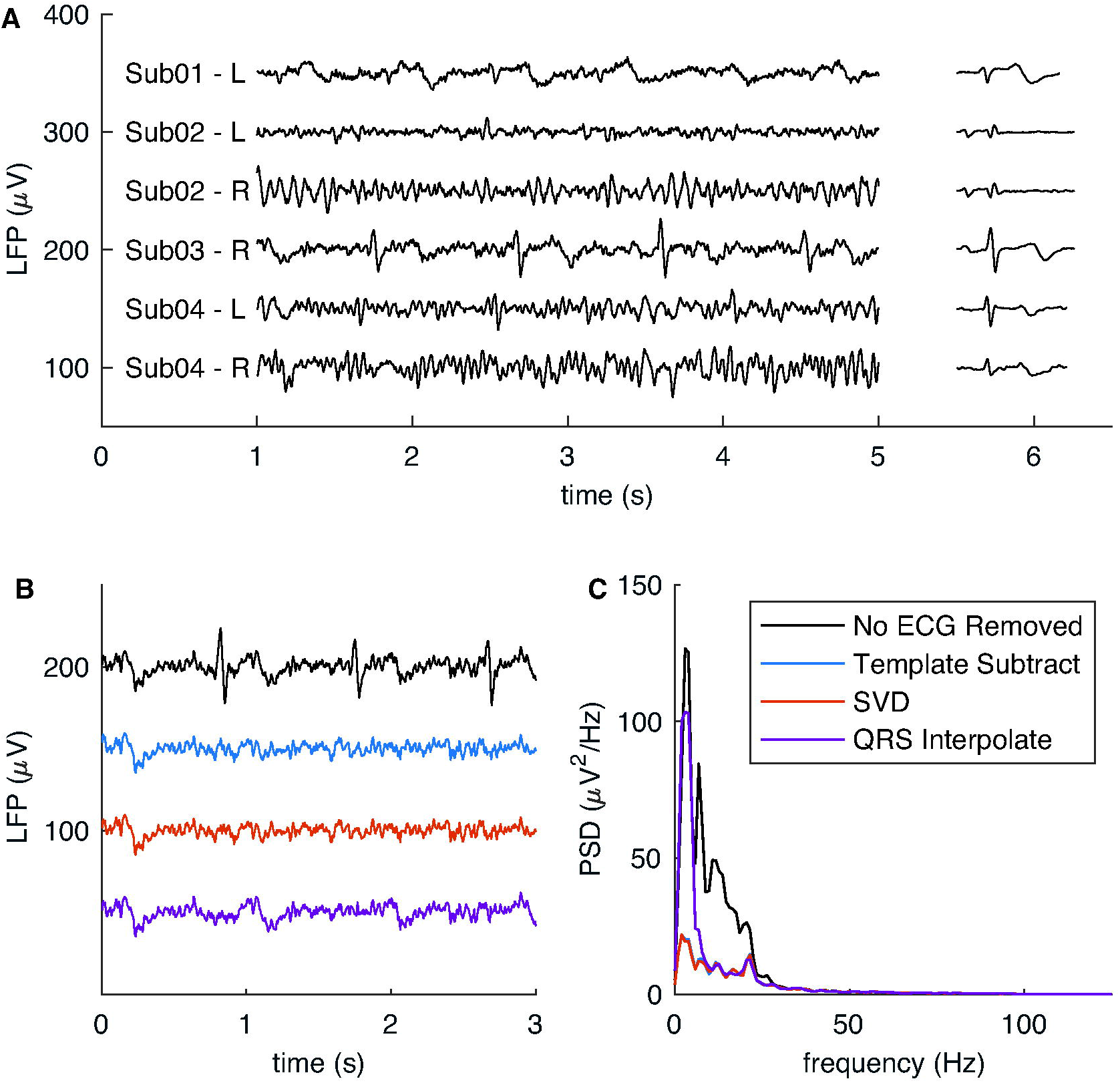
ECG artifact removal from LFP signals. (A) Sample LFP data from Sub01-Sub04 and the corresponding ECG artifact that was extracted by the template subtraction method. Comparison of ECG removal techniques and their effect on the (B) time-series data and (C) power spectrum, for a sample signal from Sub03-L. QRS interpolation left behind T-waves, which resulted in retained low-frequency content from the artifact. Both template subtraction and SVD removed this low-frequency artifact component. PSD, power spectral density.

ECG artifact morphology varied between subjects (Fig. 3A). A QRS complex was present in all cases, though the prominence of P and T waves varied. QRS interpolation resulted in adequate removal of QRS complexes, but resulted in retained low-frequency artifact in subjects with significant P or T waves (Fig. 3B-C). In contrast, both template subtraction and SVD methods removed the full ECG artifact, including both low- and high-frequency components. For each subject/hemisphere, only the first eigenvector projection calculated by SVD produced projections that extracted clear ECG artifact on visual inspection. For many of the subjects, the mean SVD projections produced a morphology very similar to the template identified by template subtraction (Fig. 4). Hemispheres that demonstrated the greatest difference between the two methods (Sub02-R and Sub04-R) were those where the first eigenvector accounted for the smallest variance explained (Table 3) and where the ECG artifact was smallest in amplitude compared to the underlying LFP signal (Fig. 3A).

**Fig. 4:**
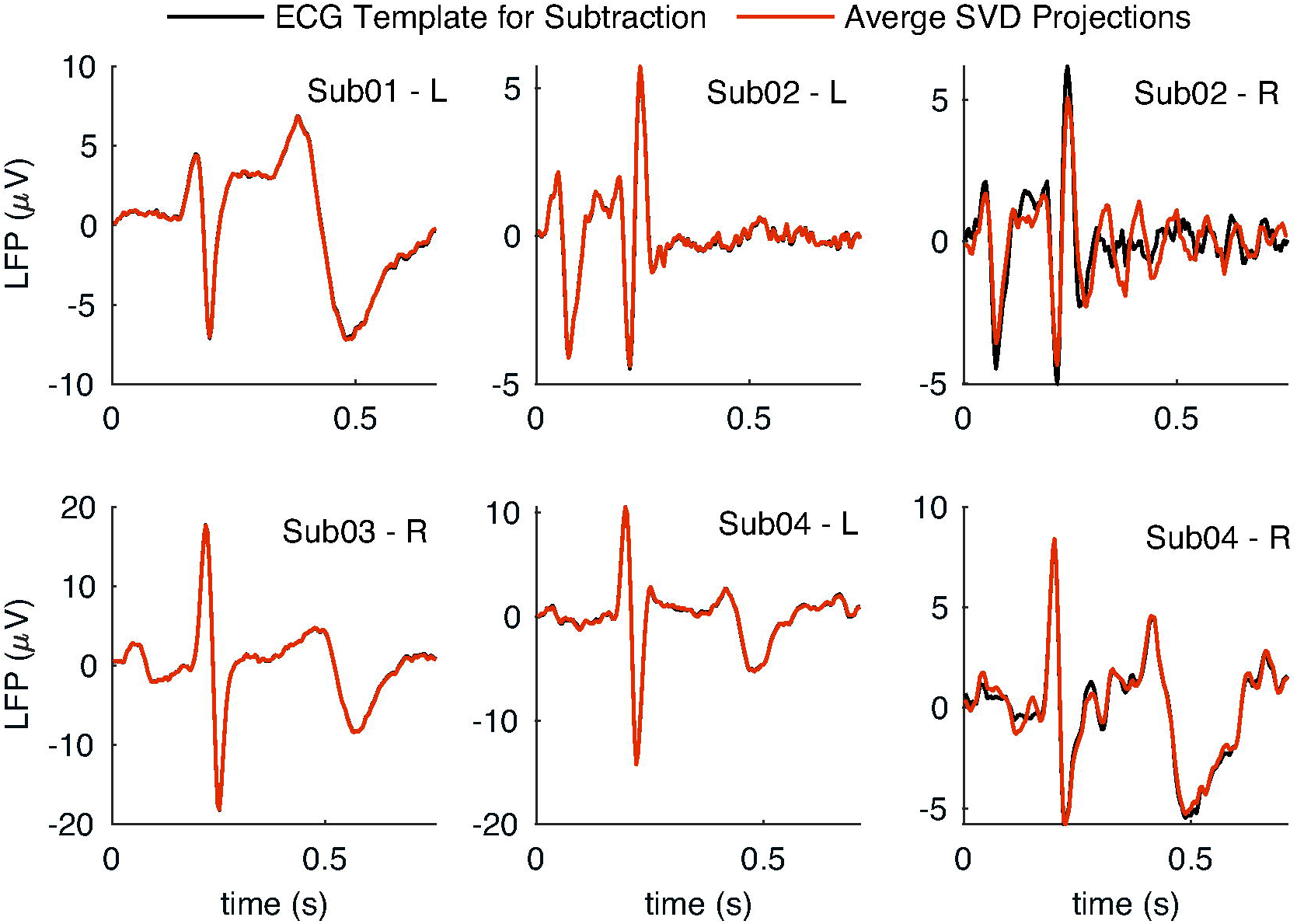
Comparison of the ECG templates used for template subtraction and averaged SVD projections for each subject. **SVD** projections include only the first principal component eigenvector. The morphology of the averaged SVD projections were similar to the template used for template subtraction. The two estimates were more likely to differ when the first eigenvector accounted for a smaller variance explained (Table 3).

**Table 3:** Variance explained by visually identified* SVD projections with appearance of ECG.

| Subject | Variance Explained |
| --- | --- |
| Sub01 – Left | 70.8% |
| Sub02 – Left | 38.4% |
| Sub02 – Right | 13.9% |
| Sub03 – Right | 75.0% |
| Sub04 – Left | 41.2% |
| Sub04 – Right | 16.4% |

\* For each subject, only the first eigenvector projection was visually identified as contributing to ECG waveform morphology

Removal of ECG artifact from ON-0mA signals by template subtraction resulted in recovery of the power spectral shape seen when stimulation was OFF (Fig. 2). SVD produced similar recovery of ON-0mA spectral shape when compared to template subtraction for all subjects and is not shown. For some subjects, although the shape was retained, the spectral magnitude was not equivalent between the two conditions, and this scaling factor was not consistent across subjects or across the two hemispheres within a subject. The presence of scaling factor differences between OFF and ON-0mA spectra required the use of spectral normalization for comparison between conditions. Averaged difference in normalized power in theta, alpha, and beta bands were less than 3.5% for both template subtraction and SVD (Table 4). QRS interpolation resulted in ON-0mA spectra that were less similar to OFF spectra. The difference between QRS interpolation and the other two methods was greatest for the theta band, which also showed the greatest variability in accuracy (89.5 ± 46.7% difference between ON-0mA and OFF for QRS interpolation).

**Table 4:** Effect of ECG removal by three different methods on the normalized spectral power in physiologically relevant frequency bands.

|  | Percentage difference of normalized power between signals recorded with stimulation ON-0mA (after artifact removal) versus stimulation OFF |  |  |
| --- | --- | --- | --- |
| Spectral Band | Template Subtraction | SVD | QRS Interpolation |
| Theta<br>(4-7.5 Hz) | -2.1 ± 7.7% | -3.4 ± 7.2% | 90.0 ± 46.6% |
| Alpha<br>(8-13 Hz) | 2.1 ± 10.5% | 0.8 ± 10.7% | 10.4 ± 16.1% |
| Beta<br>(13.5-30 Hz) | 1.7 ± 10.3% | 0.3 ± 9.9% | -4.8 ± 13.2% |

\* Mean across subjects/hemispheres ± standard error

### Non-ECG Artifact

Signals recorded within clinical care for Sub05 (STN and motor cortex electrodes) demonstrated a repetitive polyphasic artifact in the ON-0mA state (Fig. 5A). Template subtraction also successfully eliminated this artifact, removing its contribution to both low- and high-frequency content in the power spectrum (Fig. 5B).

**Figure 5:**
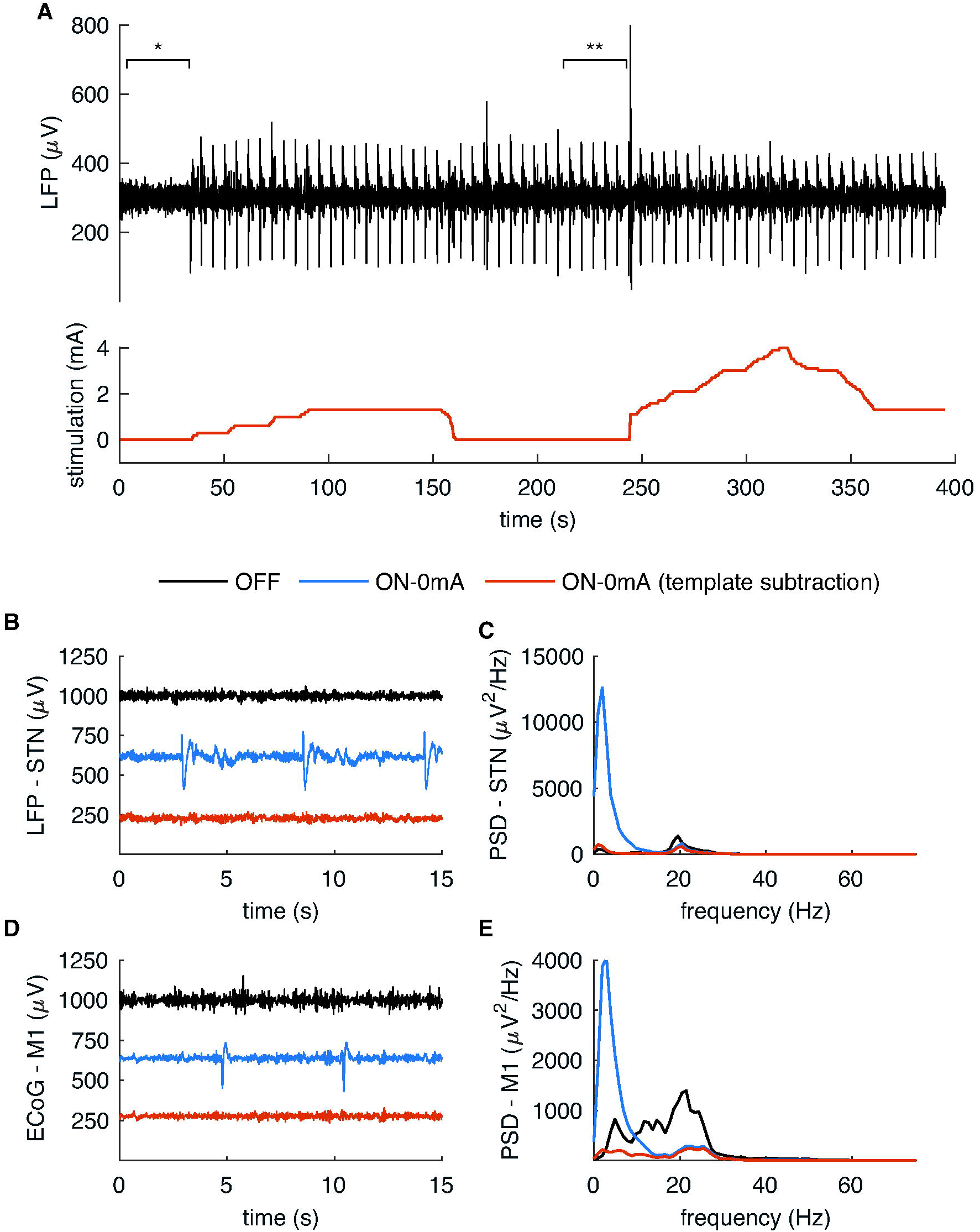
Efficacy of template subtraction on non-ECG artifact. (A) Signals collected from Sub05 as part of their standard clinical care demonstrated a repetitive, stereotyped artifact when stimulation was turned on, irrespective of stimulation amplitude. This persisted when stimulation amplitude was decreased to zero (ON-0mA). * Timepoints of data used for analysis of OFF stimulation signals in B-E. ** Timepoints of data used for analysis of ON-0mA signals in B-E. The template subtraction removed the artifact from both (B) STN and (D) M1 signals, resulting in a decrease of the low-frequency spectral content (C and E). PSD, power spectral density.

Additional sources of artifact included transient artifacts that were introduced with changes in stimulation parameters (Δ-stimulation artifact, Fig. 6). The direction of these artifacts changed depending on whether stimulation was increased or decreased, and the magnitude of the artifact increased with increasing rates of amplitude change. A transient very high-amplitude artifact was also produced in some subjects by simply enabling stimulation (turning stimulation from OFF to ON-0mA).

**Fig. 6:**
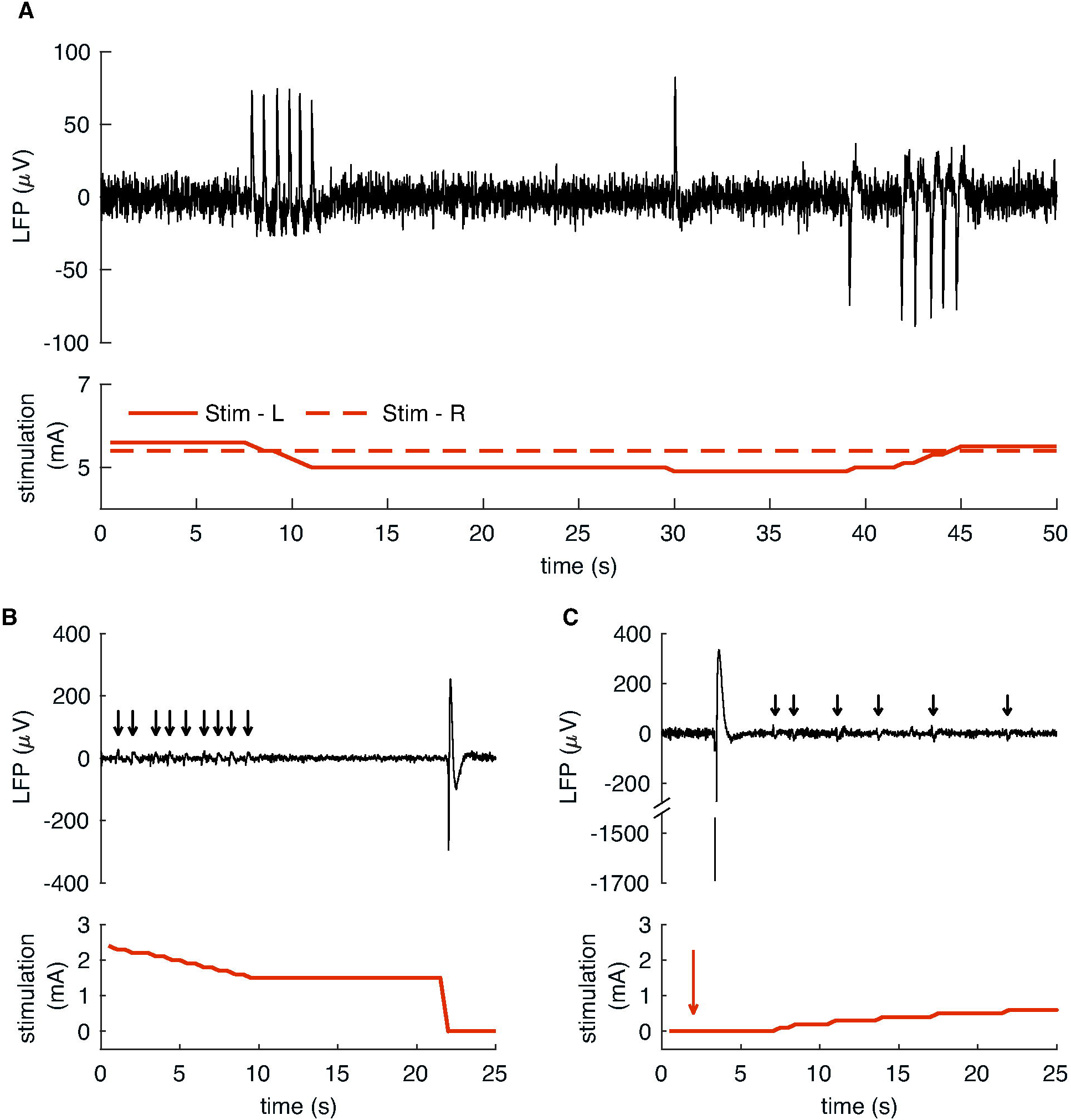
Artifact produced by changing stimulation amplitude. (A) LFP signals collected from Sub06 during stimulation ramping with concurrent Δ-stimulation artifact. (B) Signals collected from Sub07 demonstrated Δ-stimulation artifact with ramping (black arrows). A larger artifact was produced by a quicker ramp rate. (C) Additional signals collected from Sub07 again demonstrated Δ-stimulation artifact with ramping (arrows), as well as an artifact produced by switching from the OFF to the ON-0mA stimulation state (red arrow).

Signals recorded during the clinical care of Sub03 demonstrated both ECG artifact and a Δ-stimulation artifact during stimulation ramping (Fig. 7). The stimulation amplitude was ramped with a constant current increment, producing Δ-stimulation artifact with a constant magnitude and morphology. Both the Δ-stimulation and ECG artifacts were successfully removed using the template subtraction method offline. A 5 Hz frequency band centered at both 8.78 Hz (alpha range) and 16.60 Hz (beta range) of the unprocessed LFP signal showed increased power during stimulation ramping, which resolved when the Δ-stimulation artifact was removed from the signal using the template subtraction pipeline.

**Fig. 7:**
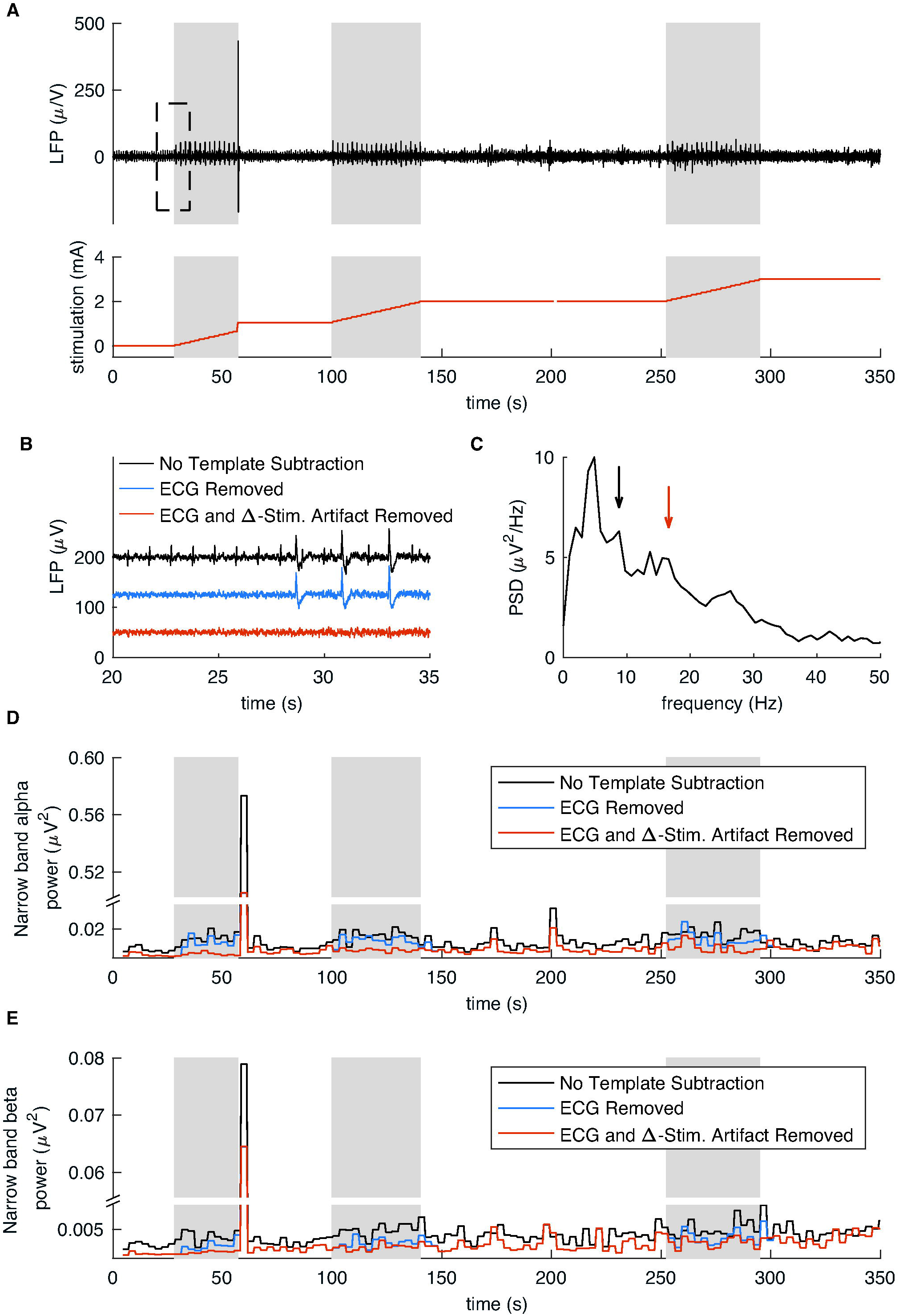
Effect of Δ-stimulation artifact on narrow-band spectral analysis. (A) LFP signals collected from Sub03 during stimulation ramping (shaded grey epochs) with concurrent Δ-stimulation artifact. Signals enclosed in the dashed rectangle are enlarged in (B), demonstrating removal of ECG and the Δ-stimulation artifact using template subtraction. (C) Power spectral density (PSD) of OFF-stimulation LFP signals demonstrated a peak at 8.57 Hz (black arrow) and 16.60 Hz (red arrow). (D) Integrated power (calculated offline) within a 5 Hz band centered on 8.57 Hz, with and without artifact removal. (E) Integrated power within a 5 Hz band centered on 16.60 Hz, with and without artifact removal.

## Discussion

This study provides an evaluation of LFP signal characteristics (including sources of artifact and potential mitigation strategies) recorded by the Medtronic Percept PC, the first commercially available sensing-enabled DBS system. We demonstrate that sensing LFP signals when stimulation is OFF is not equivalent to sensing when stimulation is ON, which has important implications for both clinical and research applications. Similar to other reports [31,32], simply enabling stimulation (even without delivered current, i.e. the ON-0mA state) introduced artifact not observed when stimulation was OFF. Sources of artifact extended beyond the electrical stimulation pulse artifact, which in contrast to previous experiences with the PC+S [21,23], did not produce subharmonic waveforms in frequency bands < 100 Hz. Artifact observed in this study included ECG (Fig. 3), a polyphasic waveform in a patient with a concurrent subdural paddle (Fig. 5), and an artifact produced during changes in stimulation amplitude (Δ-stimulation artifact, Fig. 6). These artifacts each had a stereotyped morphology, allowing them to be removed during offline post-processing with an automated template subtraction algorithm (Figs. 2-3 and 5-6). In addition to introducing artifact, enabling stimulation resulted in a multiplicative scaling difference for some patients (Fig. 2), therefore requiring normalization for within-subject comparison between OFF-vs ON-recorded signals. The presence of both artifact and scaling differences influenced spectral power estimates (Figs. 2 and 7), and therefore has the potential to affect the accuracy of LFP data used in clinical care, research, and aDBS algorithms. We recommend recording in the ON-0mA state when collecting baseline recordings aimed at biomarker detection in the absence of stimulation, both for clinical- and research-based applications. However, recording in this mode permits only a limited montage (single bipolar channel that are “stimulation-compatible”, see Table 1). If baseline richer montages are required, these can be collected OFF stimulation and processed using the pipeline enclosed herein prior to comparison with ON-stimulation signals.

ECG artifact has been similarly reported by other groups [31,32], more commonly when the IPG was implanted in the left chest wall [31]. Susceptibility to ECG, as with the PC+S, is likely due to possible insufficiency of connector seals and the use of passive recharge after stimulation, which increases the duration the electrodes are connected to the case for recharging [27]. In this study, directed analysis revealed that ECG was present in all LFP signals assessed and contributed to artifactual distortion of the PSD in lower frequency bands, even when there was no clear artifact on visual inspection of the raw time series (Figs. 2-3). Effective artifact removal techniques are therefore important to implement prior to further analysis.

Of the ECG removal methods evaluated, we chose template subtraction for rejection of both ECG and other artifacts prior to spectral analysis, given evidence of accurate artifact removal, its automated nature, and the ability to extend this method to any repetitive artifact with stereotyped morphology. Removal of ECG using template subtraction and SVD both resulted in the recovery of the original spectral pattern compared to signals recorded at baseline with stimulation OFF (with low frequency power differences of < 3.5%), suggesting near full removal of ECG from the signals (Fig. 2 and Table 4). Template subtraction assumed stationarity of the ECG artifact, which was adequate for the short-duration signals assessed in this study. A limitation of the SVD method was its visual identification of eigenvector projections. Eigenvectors were likely missed by this approach, as there may have been projections that contributed to components of the artifact, but did not resemble the full ECG artifact themselves. This limitation may explain why SVD-derived ECG artifacts differed from those extracted by template subtraction when the first principal component explained less variance (Fig. 4 and Table 3). In contrast, QRS interpolation produced less accurate recovery of power in lower frequencies, corresponding to the spectral content of P and T waves (Table 4). QRS interpolation does have the potential for quick implementation, and would be adequate for circumstances where either lower frequency bands are not of interest or where signal artifacts contain minimal P or T waves. Though a similar approach could be extended for other short-duration artifacts, the interpolation method cannot be easily extended to remove longer-duration artifacts (such as full ECG morphologies), as this would require interpolation of large percentages of the signal.

A non-ECG, polyphasic, repetitive artifact was also seen in one subject (Sub05; STN and cortical lead) when the device was in the ON state (Fig. 5). The artifact was present in multiple recordings, collected in different environments. It is unclear to what degree the connection of a subdural paddle electrode to the Percept or the presence of a contralateral Medtronic Activa SC contributed to the artifact. Template subtraction also successfully removed this artifact, providing proof of principle for removal of idiosyncratic, stereotyped artifacts. Recovery of the power spectral pattern when comparing ON-0mA versus OFF was less complete here than with the controlled experiment with Sub01-Sub04. This was likely due to the nature of the clinical recording, which lacked repetitions and randomization, and did not have a washout period to account for delayed resolution of stimulation effects [35]. The analyzed ON-0mA epoch was chosen to provide the greatest possible delay after stimulation amplitude ceased, but this may have been insufficient.

Additional artifact (Δ-stimulation artifact) occurred in many subjects when stimulation was being turned ON/OFF (similar to [32]) or the amplitude was changed (Fig. 6). Artifact did not precisely sync with recorded stimulation changes because of differences in temporal resolution between neural time series data (reported at 250 Hz), stimulation amplitude (2 Hz), and timestamps of stimulation ON/OFF changes (1 Hz). However, the artifact orientation reversed with different directions of stimulation ramping and was larger for greater rates of stimulation change, suggesting it directly resulted from the change in stimulation. Such artifacts have not been previously reported with the Percept, but have been seen with the RC+S, and were attributed to discharge of residual voltage on the electrode coupling capacitor [27]. Stimulation amplitude ramping that occurred at constant current increments produced Δ-stimulation artifact with consistent morphology and amplitude, allowing it to also be removed using offline template subtraction (Fig. 7).

The presence of a possible multiplicative scaling between OFF and ON-0mA spectra after artifact removal in some patients (Fig. 2) suggests the additional need to normalize Percept-acquired LFP data prior to further analysis. The cause of this multiplicative scaling, which was not consistent across subjects, is unclear. Reviews of the Percept’s internal hardware and sensing methods do not describe the use of different gains for ON vs OFF stimulation recordings [29,30] (confirmed directly with Medtronic). Future work should expand this evaluation to a larger subject pool to evaluate how commonly this effect is seen.

The artifact removal methods assessed were designed to be performed offline during research use, and would not be implementable in the real-time clinical setting. Onboard machine learning algorithms on the Percept flag channels with potential artifact, but their accuracy was variable. In our study, all subjects demonstrated artifact in some form, yet not all were flagged (Table 2). The clinician can override the default settings and choose to visualize the spectra of flagged channels. However, this should be done with caution as common artifacts (e.g. ECG) contain spectral content that overlap with frequency bands of clinical interest [13,18], and could confound clinical assessments of how stimulation is affecting the power spectrum. Unfortunately, the Percept system does not allow the clinician to inspect the time series signals for artifact.

A Percept-based aDBS clinical trial is underway using narrow-band beta power as a control signal [29]. Our study demonstrated that Δ-stimulation artifact transiently affected spectral content after the stimulation change (Fig. 7), which is relevant for potential aDBS implementation. The Δ-stimulation artifact produced a greater increase in spectral power at lower frequencies (theta/alpha range, corresponding to dystonia biomarkers [18] or harmonics of Parkinsonian tremor frequency [36]), though also affected narrow-band beta power (Parkinsonism [13]) as well. This artifact has the most potential to complicate single-threshold aDBS algorithms [15], where aDBS stimulation is adjusted on a shorter time-scale and may inappropriately react to artifactual increases in narrow-band power after stimulation changes. Longer timescale (i.e. dual-threshold) aDBS could also be impacted, as the frequency content of the QRS complex has the potential to confound the beta-band biomarker (particularly as cardiac rate is strongly mediated by activity levels). Future studies will be required to formally assess the effect of such artifacts on the real-time implementation of aDBS algorithms. Additionally, further studies will need to assess how the release of new DBS leads and lead extension hardware [37] affects sources of noise and potential mitigation strategies.

## Conclusion

We demonstrate that the Medtronic Percept PC’s sensing of LFP signals when stimulation is enabled is not equivalent to sensing when stimulation is OFF, because of the introduction of multiple artifacts and scaling factors. Use of (1) an automated template subtraction pipeline to remove artifacts with stereotyped waveforms and (2) spectralnormalization may allow for direct comparisons between these two conditions in an offline analysis. If not adequately removed, the presence of artifact influences estimation of spectral biomarkers.

## Statements

## Acknowledgements

We would like to acknowledge Dr. Juan Ansó for reviewing the manuscript. We also acknowledge helpful discussions with members of the OpenMind Consortium, supported by NIH BRAIN Initiative resource dissemination grant U24NS113637.

## Statement of Ethics

### Study approval statement

This study protocol was reviewed and approved by the University of California Institutional Review Board, study numbers 10-01350 and 10-02130.

### Consent to participate statement

Written informed consent was obtained from participants to participate in the study.

## Author Contributions

LHH, PAS, and SL conceived the study. LHH and RBK collected data. LHH processed and analyzed the data. LHH, PAS, and SL drafted the manuscript and figures.

## Data Availability Statement

Data collected for this study will be made available upon request.

